# Low-pressure Isochoric Freezing as a scalable technique for fresh food preservation

**DOI:** 10.1101/2024.11.01.621432

**Authors:** Alan L. Maida, Anthony N. Consiglio, Cristina Bilbao-Sainz, Andrew Karman, Gary Takeoka, Matthew J. Powell-Palm, Boris Rubinsky

## Abstract

Efficient means of reducing food waste are critically needed to meet the demands of a growing global population, while mounting consumer interest in minimally-processed food products has simultaneously driven renewed interest in non-chemical preservation modalities. Here, we investigate the potential of low-pressure isochoric freezing (LPIF) at –1.5°C / 15 MPa to simultaneously inhibit microbial growth and preserve fresh-like physiochemical and bioactive properties during storage. Using raw cow’s milk as a model product, we demonstrate synergistic effects of mild low temperature and mild enhanced pressure that significantly improve preservation for storage periods up to 5 weeks. Given the passive nature of the technique and the compatibility of the employed pressures with standard industrial compressed gas containers, these results suggest a route towards a scalable new cold storage modality.

## Main Text

The global fresh food supply chain faces the dual challenge of extending the shelf life of perishable food products while preserving their sensory and nutritional qualities. This challenge is further augmented by the increasing preference among consumers for minimal-processed foods, which, coupled with the broader goals of ensuring global food security and reducing food waste, has driven growing interest in effective non-chemical preservation techniques.

Microbial proliferation and the associated biochemical processes are often primary drivers of food spoilage^1,2^. Figure 1a illustrates various temperature and pressure-based processing and storage methods to control these factors. Generally, high temperatures and pressures are employed to inactivate microorganisms^3^, while lower temperatures are utilized to prevent their growth and the onset of spoilage reactions^1^. Often, these methods are linearly combined—e.g. deactivation treatment followed by cold storage—for greater effectiveness. While these processes delay spoilage, they often compromise nutritional and quality aspects: ice crystallization can damage cellular structures, causing textural changes and discoloration^4^; extreme conditions during high-temperature and/or high-pressure processing can denature proteins, which may reduce digestibility, degrade nutritional value, and trigger undesirable sensory changes^3^. While simultaneous application of synergistic preservation factors, like temperature and pressure, may enable the use of milder conditions, the added complexity and costs often hinder their adoption^3^.

**Figure 1.**
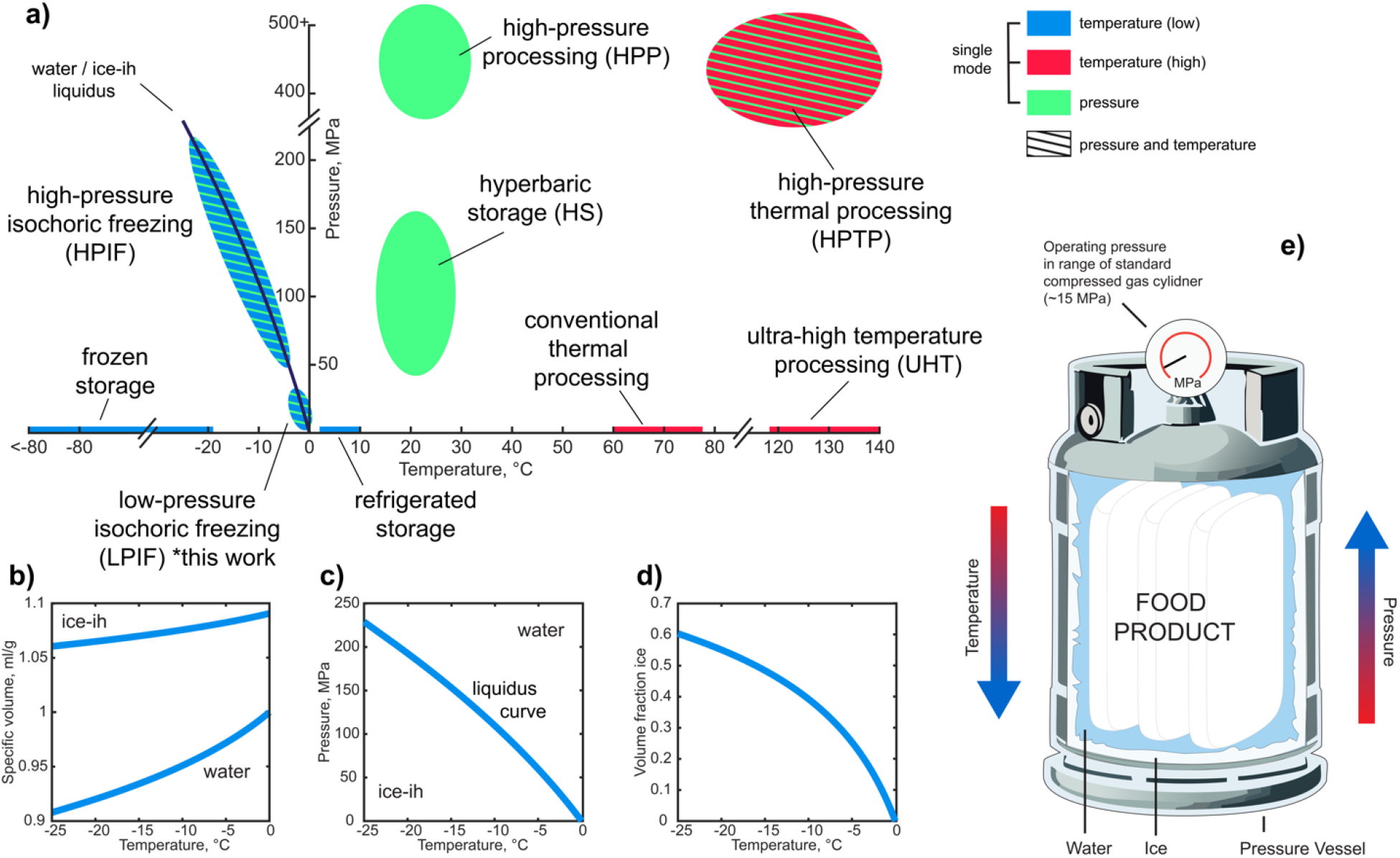
Overview of food processing/storage techniques and principles of isochoric freezing. **a)** Pressure-temperature plot depicting coordinates of various thermal- and pressure-based food processing and storage techniques. **b)** Specific volume of water and ice along the liquidus curve. **c)** Pressure-temperature water-ice liquidus curve. **d)** Ice volume fraction during isochoric freezing. **e)** Schematic of isochoric freezing preservation system with pressure monitoring. Seeded ice growth passively generates pressure while food is held in region that remains liquid.

In the past decade, a new preservation modality called isochoric freezing has attracted significant attention for its technologically simple coupling of temperature- and pressure-driven food storage processes. This technique leverages the expansion of ice-ih relative to liquid water to passively generate hydrostatic pressure within a rigid, air-free vessel during freezing, producing a two-phase liquid-ice equilibrium in which pressure and ice volume are thermodynamically coupled to temperature (Figures 1b-1d). In general, sensitive fresh food products are maintained within the unfrozen portion of the system (Figure 1e), and thereby gain the beneficial effects of temperature and pressure without suffering the consequences of direct crystallization.

Isochoric freezing has been investigated extensively for its potential to inactivate microorganisms while maintaining fresh-like nutritional and organoleptic properties in sensitive food products, with the goals of extending shelf-life and enhancing food safety^5–8^. This body of work has broadly demonstrated the synergistic effects of low temperature, enhanced pressure, and prolonged storage times, yielding significant anti-microbial outcomes at pressures often lower than the ∼50-70 MPa at which inhibition and the onset of inactivation processes are typically assumed to begin at room temperature^9–11^. However, even at the reduced pressures typical to isochoric freezing studies (∼30-90 MPa), significant concerns around the scalability of the technique remain^12,13^, given the anticipated material costs (and weight) of the thick-walled metallic vessels required to sustain these pressures. As such, studies of isochoric freezing are trending towards ever-lower pressures^14,15^, aiming to achieve sufficiently mild conditions to enable plausible industrial scalability.

To this end, we here investigate the practical low-pressure high-temperature limit of isochoric freezing (approximately –1.5°C / 15 MPa) for long term fresh food storage (up to 5 weeks). These thermodynamic conditions were selected considering that this pressure is the same encountered in myriad industrial compressed gas systems^16^, providing a key suggestion of industrial feasibility. In our goal to probe limiting behaviors, we also chose a food product, raw cow’s milk, with a notoriously active microbial population, brief shelf-life, and sensitive physiochemical profile^1^.

In order to isolate the synergistic effects of combined low temperature and mild pressure, we conducted triplicate raw milk preservation experiments to 2 and 5 weeks using low-pressure isochoric freezing (LPIF) at –1.5°C / 15 MPa, supercooled (SC) storage^17,18^ at –1.5°C / 0.1 MPa, and conventional refrigerated (RF) storage at 4°C. Each of these techniques holds the milk in a static, ice-free hypothermic state, thus enabling meaningful comparison of the effects of storage temperature and storage pressure. All experimental details are described in the Methods.

The results of this study are given in Figure 2, which shows the evolution of the microbial population, various physiochemical properties, and volatile organic compounds (VOCs) dominating flavor over a 5-week storage period under different thermodynamic conditions. LPIF at –1.5°C / 15 MPa demonstrated superior outcomes across all examined parameters, yielding a product substantially similar to the fresh control after 5 weeks. Critically, the combination of mild sub-0°C temperatures and mild enhanced pressures effectively inhibited microbial growth, yielding no significant change in the population of aerobic mesophiles over 5 weeks, and even reducing counts of the milk-dominant psychotropic (cold-resilient) pseudomonas bacteria, while both conventional refrigeration and pressure-free supercooling at –1.5°C both yielded statistically significant increases of the same. Notably, microbes are generally unaffected by such low pressures at room temperature^9,10^, further underscoring the synergistic effect of mild low temperatures and enhanced pressures on microbial inhibition.

**Figure 2.**
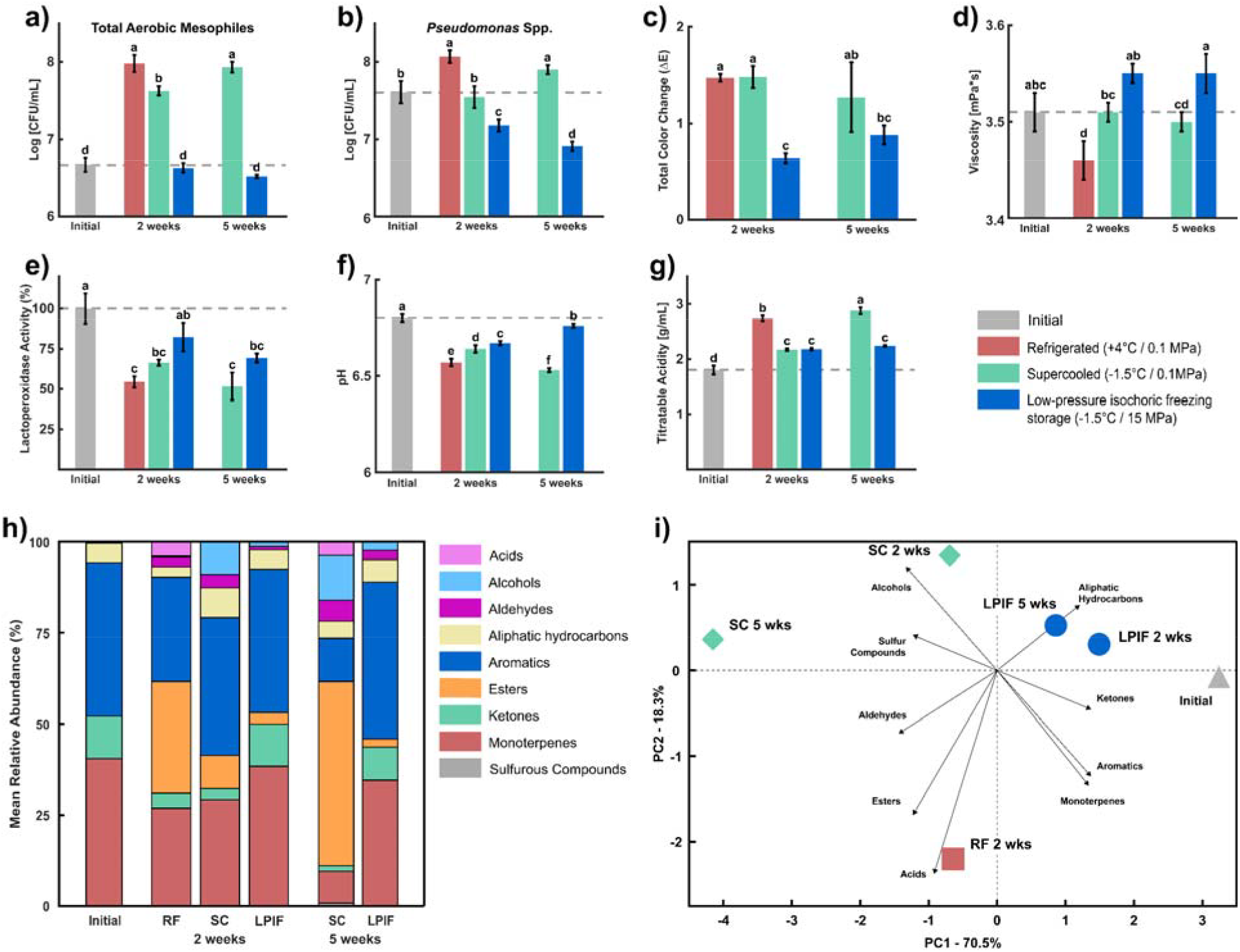
Comparison of microbial and physicochemical parameters in raw cow’s milk stored up to 5 weeks under three different conditions: refrigerated storage at +4°C / 0.1 MPa (RF), supercooled storage at –1.5°C / 0.1 MPa (SC), and low-pressure isochoric freezing storage at –1.5°C / 15 MPa (LPIF). Bar plots show (a) Total Aerobic Mesophiles (TAM), (b) Pseudomonas spp. counts, (c) Total Color Change (ΔE), (d) Viscosity, (e) pH, (f) Titratable Acidity, and (g) Lactoperoxidase Activity. Statistical difference is indicated by differing letters (p < 0.05). Error bars give standard deviation across n = 3 samples per group. Panel (h) shows the relative abundance of voaltile organic compounds evaluated using SPME-GC/MS, the raw data and statistical analyses for which are tabulated in Supplementary Table 1. PCA was also performed on this data, a biplot for which is shown in panel (i).

LPIF storage produced minimal but statistically significant changes in total color (ΔE), titratable acidity, and pH at 2 and 5 weeks (without apparent trends with time, except in the case of pH); no significant change in viscosity; and a statistically significant decrease in lactoperoxidase activity only at 5 weeks, which is notable as lactoperoxidase plays a key role in inhibiting bacterial growth and preserving freshness in milk^18^. Importantly, LPIF samples at – 1.5°C and 15 MPa experienced lesser deviations from control across all parameters and time points.

We also characterized the volatile organic compounds (VOCs) profile of each experimental group using solid-microextraction coupled with gas chromatography/mass spectrometry (SPME-GC/MS), identifying 28 volatile compounds across nine classes. All compounds and concentrations are listed in Supplementary Table 1 with statistically significant changes indicated, and the relative abundance of each class of compounds is shown in Fig. 2h for each experimental condition and time point. Principal components analysis (PCA) was used to further elucidate the evolution of the VOCs profile across groups, a biplot for which is shown in Fig. 2i. Both figures indicate substantial consistency in the flavor profile of the LPIF group as compared to control, especially with respect to the avoidance of esters (e.g. ethyl acetate) associated with spoilage and fermentation defects^19^, which dominate refrigerated and supercooled samples at 5 weeks, and the preservation of the native aromatics profile, which does not significantly differ from control over entire storage period.

Collectively, these results suggest that the combination of mild sub-zero temperatures (– 1.5°C) and mild enhanced pressures (15 MPa) provides a thermodynamic environment that broadly inhibits spoilage bacteria and arrests deleterious biochemical processes, thereby preserving the physicochemical integrity of sensitive fresh food products. While our study is limited to raw cow’s milk, we note that this model provides a uniquely rich bacterial and biochemical template. In light of this fact, the widely recognized role of *Pseudomonas* spp., in spoilage of animal-based products and common refrigerated foods^1,2^, and the wide range of products over which similar results have been observed in higher-pressure isochoric freezing protocols, we suggest that these conditions may prove broadly applicable to prolonged preservation of foods in a fresh-like, microbially-inhibited state.

Finally, we note that the pressurization of isochoric freezing systems is driven solely by temperature, requiring no moving parts or active mechanical processes. Given that the pressures employed align with those used in industrial compressed gas cylinders, we suggest that low-pressure isochoric freezing may provide a simple and scalable technology for prolonged storage of fresh food products, which may be employed to reduce post-harvest food waste, aid diffusion of these products into current food deserts, and increase global food capacitance and security.

## Acknowledgements

This work was supported by the USDA National Institute of Food and Agriculture, AFRI project Proposal #:2021-09570, Award #2022-67017-37098 “Novel Isochoric Processing for Sustainable, Safe and High-Quality Preservation of Fluid Foods”. A.M. also recognizes support from the National Science Foundation Graduate Research Fellowship Program under Grant No. DGE 2146752.

## Data availability

All data are available upon reasonable request to the corresponding author.

## Competing interests

A.N.C, M.J.P.P., and B.R. have a financial stake in BioChoric Inc., a private entity working on commercialization of isochoric freezing technologies. The rest of the authors declare no competing interests.

## Materials and Methods

### 1.1. Isochoric Freezing System

Raw milk was treated at –1.5 °C / 15 MPa using a pressure chamber made of Aluminum-7075 with a type-II anodize coating and a total volume capacity of 1500 mL, pressure-rated for up to 275MPa (BioChoric Inc., Bozeman, MT, USA). Each chamber was connected to a pressure gauge to monitor the pressure over time. The chambers were cooled using a chest freezer (Magic Chef Model #HMCF9W3, *MC Appliance Corporation*, Wood Dale, IL, USA).

### 1.2. Supercooled System

Raw cow milk was placed into 15 mL sterile conical centrifuge tubes made of polypropylene (Heathrow Scientific). To ensure stability of the supercooled state over an extended period, each tube was completely filled to eliminate any air interface, effectively mimicking isochoric conditions for enhanced supercooling stability. The absence of air ensured that any freezing would cause visible expansion of the container, providing an immediate indication of ice formation. The clear polypropylene material also allowed for easy visual detection of any supercooling failure. This method enabled consistent monitoring and validation of the supercooled state throughout the 5-week storage period. Cooling was achieved using a chest freezer (Magic Chef Model #HMCF9W3, MC Appliance Corporation, Wood Dale, IL, USA)

### 1.3. Experimental protocol

Raw bovine milk was purchased from a local market located in Albany, California. Three different conditions were compared: conventional refrigeration (RF) at 4°C, supercooled (SC) preservation at –1.5 °C / 0.1 MPa, and low-pressure isochoric freezing (LPIF) preservation at – 1.5°C / 15 MPa. Initial tests were conducted seven days before the labeled expiration date of the milk. The initial quality of the unprocessed raw cow’s milk was immediately assessed to measure microbiological counts, and physiochemical properties. All the treatment samples of the milk were prepared on the same day, with subsequent tests conducted after 2 and 5 weeks of storage.

For LPIF, approximately 300 mL of raw cow’s milk was packed into two sterilized polyethylene bags (Whirl-Pak®). The bags were heat sealed with negligible air bubbles and transferred to two separate isochoric chambers (total of four bags) filled with cool water. After the treatment period, the chambers were removed from the freezers and placed in a refrigerator set at 4 °C for 12h to allow all ice within the chamber to melt and the pressure to recede accordingly.

For SC conditions, a total of 40 tubes were used, with 30 designated for the experiments and 10 reserved as backups in case of freezing or other complications. After the treatment, the supercooled samples were removed from the freezers and placed in a refrigerator set at 5 °C for one hour to warm above the freezing temperature.

### 1.4. Microbiology

Decimal dilutions were prepared for total plate count and *Pseudomonas*. Appropriate dilutions were plated on plate count agar (PCA) for total aerobic mesophiles (TAM) and incubated at 30°C for 3 days (ISO 4833:2013).

Numbers of *Pseudomonas* spp. were determined by spread plating appropriate dilutions on *Pseudomonas* agar base with Cephalothin, Fucidin, Cetrimide (CFC) supplement (Oxoid Ltd., Basingstoke, Hampshire, U.K.) and incubated at 25°C for 72 hours.

### 1.4. pH/Titration

The pH of the milk was determined in triplicate using a pH meter (Hanna instruments, USA). The titratable acidity (TA) of the milk samples was determined by titrating 10 mL of diluted milk (4 mL of milk in 6 mL of DI water) to pH of 8.4 with 0.01 M sodium hydroxide solution. The results were expressed as grams of lactic acid per liter of milk based on:

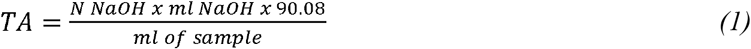

### 1.5. Color

The color of the bovine milk was measured using a tristimulus colorimeter (CM508D, Konica-Minolta Inc., Ramsey, NJ, USA) with a sample holder (CM-A128) and an 8 mm diameter target mask (CM-A195). The milk sample (10 mL) was pipetted into the sample holder covered with the black background and the color for each sample was measured three times. Results were expressed as L* (lightness), a* (redness/greenness), and b* (yellowness/ blueness) in the CIE Lab system. These values were used to calculate the overall color difference (ΔE*) according to the following equations:

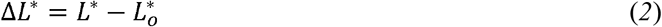

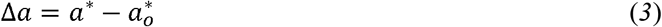

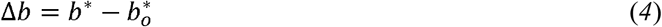

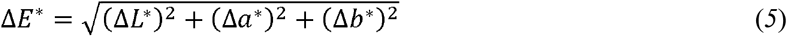

where L_o_^*^, a_o_^*^, and b_o_^*^ represent the initial values prior to storage/treatment at day 0.

### 1.7. Viscosity

Milk viscosity measurements were performed on a DHR-3 rheometer (TA Instrument, New Castle, DE) at 20°C. A concentric cylinder geometry (28.04 mm bob diameter and 21.10 mm bob length) was used for testing. Three replicates were performed for each sample. For each replicate, 8 mL of milk was pipetted into the concentric cylinder cup and equilibrated at 20°C for three minutes. The shear rate was increased logarithmically from 0.1-200 s^−1^. The viscosity of the sample was determined at a shear rate of 100 s^−1^.

### 1.8. Lactoperoxidase Activity

The lactoperoxidase (LPO) activity assay was performed based on the method described by Maida et al. ^6^. Briefly, milk was mixed with a solution of 0.65 mM ABTS (in 0.1 M sodium phosphate bu□er, pH 6.0) and left for 30 min at 20 °C. Then, 0.1 mM hydrogen peroxide was added and mixed quickly to initiate the reaction, with the absorbance (Abs_412nm_) measured for 1 minute. The enzymatic activity was calculated as the slope of Abs increment as a function of time and expressed as ΔAbs_412nm_ AU/min. All enzymatic assays were performed in triplicate for each storage condition. The residual activity was calculated by using:

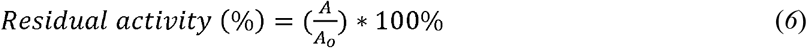

where *A* is the enzymatic activity in milk samples after storage and *A*_0_ represents the initial enzymatic activity prior to storage.

### 1.9. HS–SPME–GC/MS Analysis of Volatiles

Volatiles were analyzed in accordance with the headspace solid-phase microextraction gas chromatography mass spectrometry (HS**–**SPME–GC/MS) method reported in Maida et al (2023). First, a 250 mg/L internal standard solution was prepared by dissolving cyclohexanone (> 99 %, MilliporeSigma, Burlington, MA) in deionized water (18 MΩ·cm). Next, a 10 mL aliquot of raw milk, 0.74 g of sodium chloride, 250 µL of internal standard solution, and a magnetic stir bar was added to a 240 mL amber screw vial. Each vial was immediately sealed together with a 20 mm PTFE/silicone-lined septum (MilliporeSigma) and a plastic screw cap (Sigma Aldrich, St. Louis, MO) combination. Afterwards, the vials were held at 50°C in a heated water bath for 15 minutes under mild stirring to reach temperature equilibration. Temperatures were monitored using a calibrated digital thermometer (Anritsu, Atsugi, Japan). HS**–**SPME sampling was performed using a 2-cm polydimethylsilioxane/divinylbenzene/carboxen (PDMS/DVB/CAR, Supleco, Bellefonte, PA) 23-gauge SPME fiber. Following temperature equilibration, the SPME fiber was exposed to the headspace of solution under the same conditions for 30 min prior to analysis by GC/MS.

After headspace extraction, the fiber was desorbed for 10 min in the injection port (250°C) of a gas chromatograph (7890B) coupled to a mass selective detector (5977A; Agilent Technologies, Santa Clara, CA). A DB-1 fused silica capillary column (60 m × 0.25 mm d_f_ = 0.25 µm, J&W Scientific, Folsom, CA) was used together with ultra-high purity helium gas (99.999%, Matheson, Richmond, CA) that was held at a constant flow rate of 1 mL/min. The GC oven program was as follows: 35°C for 5 minutes, ramped at 4°C/min until 100°C, a second ramp 10°C/min until 220°C, and finally held at 220°C for 5 minutes, with a total run time of 41.25 minutes. The MS source was set at 230°C, the quadrupole at 150°C, and ionization energy at 70eV. The MS was run in full scan mode at *m/z* 20 to 350 at 2.3 scans/s. The purge-to-split valve was opened after 1 min at a flowrate of 30mL/min following sample injection.

Data processing was performed using MassHunter Unknowns Analysis (v. 10.1, Agilent Technologies, Santa Clara, CA). Parameters used for deconvolution consisted of 0.3 for left *m/z* and 0.7 for right *m/z* delta extraction window and an RT size window of 100. Retention indices (RI) were calculated from the analysis of a calibration file obtained from an alkane mixture (C_5_-C_18_, Sigma Aldrich, St. Louis, MO, USA). Compounds putatively identified by the mass spectra were limited to only those with a match score of ≥70% and an ion signal to noise ratio ≥ 10. In addition, the calculated RI were compared to those previously reported in the literature. Analytes associated with column bleed (i.e., silanes and siloxanes) or artifacts from the fiber were discarded. Quantitation of volatiles was performed by taking the ratio of each volatile compound peak area to the internal standard peak area.

### 1.10. Statistics

All experiments and analyses were carried out in triplicate. Analysis of variance (ANOVA) was performed under all the different storage conditions, followed by a multiple comparison post hoc test, Tukey’s HSD test, at a 5% level of significance. Additionally, principal component analysis (PCA) was performed to identify the statistical patterns in VOC data.

