## Supplementary Table 1 for "Low-pressure Isochoric Freezing as a scalable technique for fresh food preservation"

**This PDF file includes:**

Supplementary Table S1

**Table S1**. Volatiles expressed as µg equivalent internal standard per 100 mL of milk. Different letters across a row denote significant differences at $p < 0.05$.

| Class | Compound | $RI_r$[1] | $RI_e$ | Peak[2] | I[3] | RF[3] | LPIF[3] | LPIF[3] | SC[3] | SC[3] |
|---|---|---|---|---|---|---|---|---|---|---|
| | | | | | 0[4] | 2[4] | 2[4] | 5[4] | 2[4] | 5[4] |
| Acids | Octanoic acid | 1165 | 1156 | 25 | ND | 4.00 a | ND | ND | ND | 3.07 a |
| | n-Decanoic acid | 1335 | 1344 | 28 | ND | 0.97 | ND | ND | ND | ND |
| Sum | | | | | ND | 4.97 a | ND | ND | ND | 3.07 a |
| Alcohols | Ethanol | 440 | 466 | 1 | ND | ND | ND | ND | 3.06 a | 5.65 a |
| | 1-Butanol, 2-methyl- | 729 | 723 | 5 | ND | ND | ND | ND | ND | 1.85 |
| | 1-Hexanol | 852 | 854 | 11 | ND | 0.42 d | 1.13 c | 1.81 b | 1.41 bc | 2.56 a |
| Sum | | | | | ND | 0.42 c | 1.13 c | 1.81 c | 4.46 b | 10.06 a |
| Aldehydes | Hexanal | 783 | 775 | 6 | ND | 1.50 | ND | ND | ND | ND |
| | Nonanal | 1082 | 1084 | 24 | 0.47 b | 1.69 ab | 0.79 b | 1.96 ab | 1.79 ab | 4.66 a |
| Sum | | | | | 0.47 c | 3.19a | 0.79 bc | 1.96 b | 1.79 b | 4.66 a |
| Aliphatic hydrocarbons | Octane | 800 | 800 | 8 | ND | ND | ND | 0.54 a | 0.6 a | 0.48 a |
| | Heptane, 2,4-dimethyl- | 824 | 824 | 9 | 1.12 a | ND | 1.17 a | 0.64 a | 1.22 a | 1.57 a |
| | R(-)3,7-Dimethyl-1,6-octadiene | 926 | 945 | 15 | 3.56 a | 2.11 b | 2.25 b | 1.96 bc | 0.87 cd | 0.28 d |
| | Heptane, 2,2,4,6,6-pentamethyl- | 1003 | 1003 | 18 | 1.30 a | 1.45 a | 1.02 a | 1.52 a | 1.38 a | 1.51 a |
| Sum | | | | | 5.98 a | 3.56 ab | 4.44 ab | 4.66 ab | 4.07 b | 3.84 b |
| Aromatics | *p*-Cymene | 1014 | 1015 | 19 | 46.7 a | 34.99 a | 32.84 ab | 32.45 ab | 18.15 bc | 9.04 c |
| | Benzene, 1-methyl-3-(1-methylethenyl)- | 1075 | 1077 | 22 | 0.71a | 0.71 a | 0.41ab | 0.35 ab | 0.32 b | 0.32 b |
| | Benzene, 1,3-bis(1,1-dimethylethyl)- | 1247 | 1250 | 27 | ND | ND | ND | ND | 0.25 a | 0.34 a |
| Sum | | | | | 47.41 a | 35.7 ab | 33.25 ab | 32.80 b | 18.72 c | 9.70 d |
| Esters | Ethyl Acetate | 601 | 601 | 4 | ND | 14.44 b | 2.66 c | 1.70 c | 4.52 bc | 35.41 a |
| | Butanoic acid, ethyl ester | 793 | 785 | 7 | ND | 8.00 a | ND | ND | ND | 5.85 a |
| | Butanoic acid, 3-methyl-, ethyl ester | 844 | 838 | 10 | ND | 2.51 | ND | ND | ND | ND |
| | Hexanoic acid, ethyl ester | 986 | 983 | 17 | ND | 11.53 | ND | ND | ND | ND |
| | Octanoic acid, ethyl ester | 1184 | 1180 | 26 | ND | 1.68 | ND | ND | ND | ND |
| Sum | | | | | ND | 38.16 a | 2.66 b | 1.70 b | 4.52 b | 41.26 a |
| Ketones | Acetone | 468 | 486 | 2 | 8.78 a | 3.30 c | 6.78 b | 5.57 b | 0.65 d | 0.83 d |
| | 2-Butanone | 575 | 570 | 3 | 4.28 a | 1.82 c | 2.99 b | 1.38 cd | 0.86 de | 0.43 e |
| Sum | | | | | 13.06 a | 5.12 c | 9.77 b | 6.95 c | 1.51 d | 1.26 d |
| Monoterpenes[5] | α-Phellandrene | 995 | 926 | 13 | 0.48 a | 0.30 b | 0.29 b | 0.26 b | ND | ND |
| | Bicyclo[3.1.1]hept-2-ene, 3,6,6-trimethyl- | 948 | 933 | 14 | 2.28 a | ND | 1.74 b | ND | ND | ND |
| | ß-Pinene | 973 | 974 | 16 | 4.76 ab | 5.94 a | 3.72 ab | 3.63 ab | 0.43b | 1.1 ab |
| | D-Limonene | 1023 | 1026 | 20 | 31.19a | 22.50a | 22.05ab | 21.39ab | 11.56 bc | 5.18 c |
| | 3-*p*-Menthene | 976 | 1029 | 21 | 5.66 a | 3.97 a | 3.9 a | 0.26 b | 1.97 b | 0.83 b |
| | δ-Carene | 1025 | 1083 | 23 | 1.22 a | 0.82 ab | 0.74 b | 0.78 b | 0.53 b | ND |
| Sum | | | | | 45.6 a | 33.52 ab | 32.43 bc | 26.32 c | 14.49 d | 7.11 e |
| Sulfur Compounds | 2,4-Dithiapentane | 873 | 865 | 12 | ND | ND | ND | ND | ND | 0.70 |

[1]Reference retention index obtained from http://webbook.nist.gov (accessed 05/29/2024) or the NIST MS library (2014). [2]Peaks numbered by earliest elution time. [3]Treatment types: I = initial, RF = refrigeration, LPIF =low-pressure isochoric freezing (-1.5°C / 15 MPa), SC = super cooling (-1.5°C / 0.1MPa)[4] 0,2,5 refer to number of weeks in storage for a given treatment. [5]Monoterpenes and modified monoterpenes have been grouped together.
